# Heterogeneity and Metabolic Diversity among *Enterococcus* Species during Long-term Colonization

**DOI:** 10.1101/2024.10.18.619042

**Authors:** Philip A. Karlsson, Taoran Zhang, Josef D. Järhult, Enrique Joffré, Helen Wang

**Affiliations:** Department of Medical Biochemistry and Microbiology, Uppsala University, Uppsala, Sweden; Department of Medical Sciences, Zoonosis Science Center, Uppsala University, Uppsala, Sweden; Department of Microbiology, Tumor and Cell Biology, Stockholm, Sweden; Department of Chemistry and Molecular Biology (CMB), University of Gothenburg, Gothenburg, Sweden

**Author notes:** Corresponding author: **Helen Wang (**). Philip A. Karlsson and Taoran Zhang contributed equally to this work. Author order was determined on the basis of seniority.

## Abstract

Urinary tract infections (UTIs), traditionally dominated by Gram-negative pathogens, are increasingly complicated by antimicrobial-resistant *Enterococcus* spp. in hospital settings. This study screened urine samples from 210 ICU patients at Uppsala University Hospital (June 2020 - September 2021), identifying 39 unique PhenePlate™-RF types across *E. faecium*, *E. faecalis*, and *E. durans*. *E. faecium* isolates showed considerable genetic diversity, primarily within clonal complex 17 (CC17), known for its virulence and antibiotic resistance. We identified multiple lineages and sequence types (STs), such as in patient HWP143, who had isolates from both ST80 and ST22 (an ancestral CC17 lineage). Notably, metabolic adaptations, such as increased L-arabinose metabolism, and shifts in antibiotic resistance were observed. Variations and similarities in plasmid content between individual linages suggest horizontal gene transfer. *E. faecalis* isolates exhibited less genetic diversity but significant metabolic variability across patients and mixed infections, as seen in patient HWP051, colonized by both ST16 (CC58) and ST287. *E. durans*, though less common, shared important metabolic traits with *E. faecium* and displayed polyclonal characteristics, highlighting its potential role in UTIs and the complexity of enterococcal infections. *E. durans* was sometimes misidentified, underscoring the need for accurate identification methods. This research underscores the importance of understanding genetic and metabolic diversity, plasmid variations, and horizontal gene transfer in *Enterococcus* spp., which influence antibiotic resistance, virulence, and ultimately, treatment outcomes.

**IMPORTANCE STATEMENT:** Our study uncovers novel insights into the genetic and metabolic diversity of *Enterococcus* species within individual patients, focusing on *E. faecium*, *E. faecalis*, and *E. durans*. Unlike prior studies, which often focused on single lineages, we reveal multiple clones and lineages across individual patients, including clones from clonal complex 17 and the emerging sequence type (ST) 192, highlighting notable metabolic adaptations and shifts in antibiotic resistance. The detection of mixed colonization with varied ST-types, and *E. durans* misidentification by MALDI-TOF, later corrected by sequencing, further emphasizes the challenges in *Enterococcus* species identification. For the first time, we demonstrate likely horizontal gene transfer among *E. faecium*, *E. faecalis*, and *E. durans* within the same patient, underscoring the dynamic nature of these infections. Our findings have significant implications for understanding the complexity of *Enterococcus* infections, stressing the need to consider genetic and metabolic diversity to improve disease management and treatment outcomes.

## INTRODUCTION

Urinary tract infections (UTIs) are among the most prevalent bacterial diseases, posing a considerable burden to public health, particularly in the face of rising antimicrobial resistance (1). Risk factors of UTIs include pregnancy, anatomical and functional abnormalities of the urinary tract, and the use of indwelling urinary catheters (2).

Gram-negative pathogens have traditionally been considered the primary causative agents of UTIs (3), however, recent studies reveal an increasing isolation of *Enterococcus* species in bladder infections (4, 5). Historically, Uropathogenic *Escherichia coli* (UPEC) has been responsible for 75-85% of all UTIs. Although Gram-positive bacteria constitute less than 15% of community acquired UTIs, they are more frequently associated with nosocomial infections (6, 7). *Enterococcus*, naturally resistant to many first-line antibiotics and capable of forming biofilms, contributes to immune evasion and treatment failure (8). Certain *Enterococcus* strains can acquire high-level resistance to ß-lactams, aminoglycosides, glycopeptides, and even combined antibiotic therapies. In Scandinavia, vancomycin-resistant *Enterococcus* are rarely found but are causing outbreaks at an increasing and alarming speed (9–11). Ampicillin-resistant *E. faecium* are gradually more common in hospital-acquired infections (12). In the United States, *Enterococcus* spp. were responsible for 12% of all catheter-associated UTIs (CAUTI) between 2006 and 2007 (13). This is likely due to their ability to elicit proinflammatory responses in the bladder and form biofilm (14), which enable them to persist and cause chronic infections inside the bladder.

Bacterial populations are inherently heterogeneous, providing selective advantages during environmental changes, and profoundly influencing clinical outcomes (15). Recent studies have demonstrated that *E. faecalis* exhibits heterogeneity in adhesion and biofilm formation. These key virulence properties may contribute to prolonged hospital stays and treatment failure (16).

In this study, we focus on bacterial urine colonizers among critically ill patients with severe acute respiratory syndrome coronavirus 2 (SARS-CoV-2) infection. Early studies from similar cohorts have revealed that severely ill patients have a significantly higher risk of acquiring bacterial co-infections, especially with resistant strains, due to factors such as increased antibiotic dosage, utilization of urinary catheters, and the administration of immunosuppressive drugs (17, 18). A 2022 study in Madrid investigated 87 COVID-19 patients, 89.6% had acquired UTIs, with 67.9% being related to CAUTIs. *Enterococcus* was identified as dominant species, representing 47.4% of the study cohort (4). However, the role of heterogeneity and dynamics in these conditions remain unknown.

Distinguishing between species and clinically relevant strains of *Enterococcus* can be challenging, and their rapid emergence and importance demand a fast and accurate screening method (19, 20). In this study, we utilized the PhenePlate™-RF (PhP-RF) system, a method for strain screening that provides a biochemical fingerprint for multiple clinical *Enterococcus* species, first introduced by Kühn et al. (21). This method is fast, highly reproducible, and possesses strong discriminatory power. The PhP-RF system has been successfully used and validated for screening *Enterococcus* in multiple environments, including the food chain and sewage water (22). However, its application to clinical *Enterococcus* isolates has been rare (17). Our study employs the PhP-RF system to investigate the heterogeneity and dynamics of clinical *Enterococcus* during bladder colonization.

## RESULTS

From June 2020 to September 2021, urine samples were collected from 210 patients undergoing intensive care at Uppsala University Hospital. A total of 29 urine samples, pre-determined based on *Enterococcus* presence, were screened using the PhP-RF system. In total, 456 colonies (*E. faecium*: n=245, *E. faecalis*: n=194, *E. durans*: n=17) were assessed, and 39 PhP-types (*E. faecium*: n=25, *E. faecalis*: n=8, *E. durans*: n=6) were identified using the UPGMA clustering approach. One representative isolate per PhP-type and patient isolation day was confirmed with MALDI-TOF and WGS (Supplementary Figure 1).

### *Enterococcus faecium* isolates showed high heterogeneity

*E. faecium* isolates were grouped into two major PhP-branches, seemingly related to raffinose metabolism. The first branch included all strains from patient HWP004 and HWP199, and two isolates from patient HWP143 (**Figure 1A**). The third strain from patient HWP143 did not group with either of these branches, standing alone in the tree. This separation among HWP143 isolates was also observed genetically, collectively indicating heterogeneity (**Figure 1B**). This genetic separation is further supported by the isolates carrying several different plasmids (Supplementary Figure 2). Isolate HWP004:11E could not have its ST-type determined through the chosen method, but distinctly grouped among ST127 isolates, potentially suggesting a new ST variant or mixed sample. All HWP004 isolates belonged to different PhP-types, yet HWP004:11E was the only HWP004-isolate grouping differently phylogenetically. The different PhP-types across the different days within one patient were presented as proportions of subpopulations (**Figure 1C-D**). HWP004:11E was the only HWP004 strain demonstrating resistance to tobramycin (TOB), and did not carry the repUS15 plasmid, which was present in all other HWP004 strains (**Figure 2, Supplementary Figure 2**). Instead, HWP004:11E carried the rep1 plasmid, which was lost in two isolates on collection day 12 (HWP004:12A and HWP004:12F) but was omnipresent during collection day 13 (suggestively through isolate HWP004:12B). Unlike all other HWP004 strains, HWP004:12A did not tolerate TZP.

**Figure 1.**
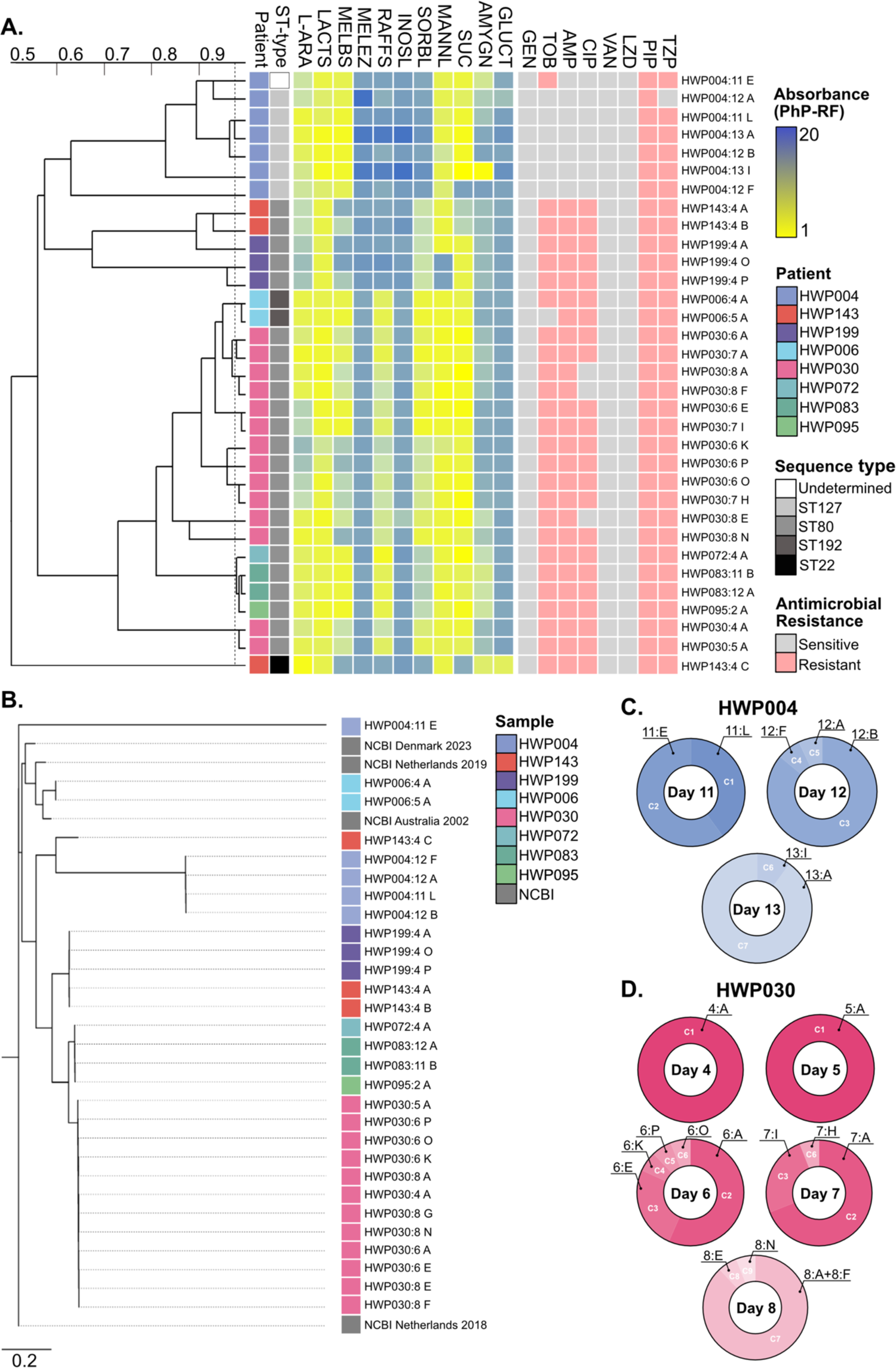
Heterogeneity of *E. faecium* in the patient cohort. **A.** Dendrogram derived from Unweighted Pair Group Method with Arithmetic Mean (UPGMA) clustering of PhP-RF data (absorbance values at 620nm). The blue-yellow gradient represents a scale from low (blue) to high (yellow) metabolism. Isolates with the most recent branching point above the 0.975 cut-off value (dotted line) are grouped into a PhP-type. For a full list of metabolic reagents, refer to the material and methods or supplementary data. Phenotypic resistance (pink) and susceptibility (gray) were assessed using broth microdilutions according to EUCAST. ST-types were determined based on Illumina WGS. **B.** Phylogenetic tree based on Single Nucleotide Polymorphism (SNP) comparison of Illumina WGS data. Three NCBI strains are included as comparative references based on diversified historical and geographic background. **C.** PhP-type cluster diversity (proportion) based on the number of colonies from the total number screened (n=16) for isolates from patient HWP004. **D.** PhP-type cluster diversity (proportion) based on the number of colonies from the total number screened (n=16) for isolates from patient HWP030.

**Figure 2.**
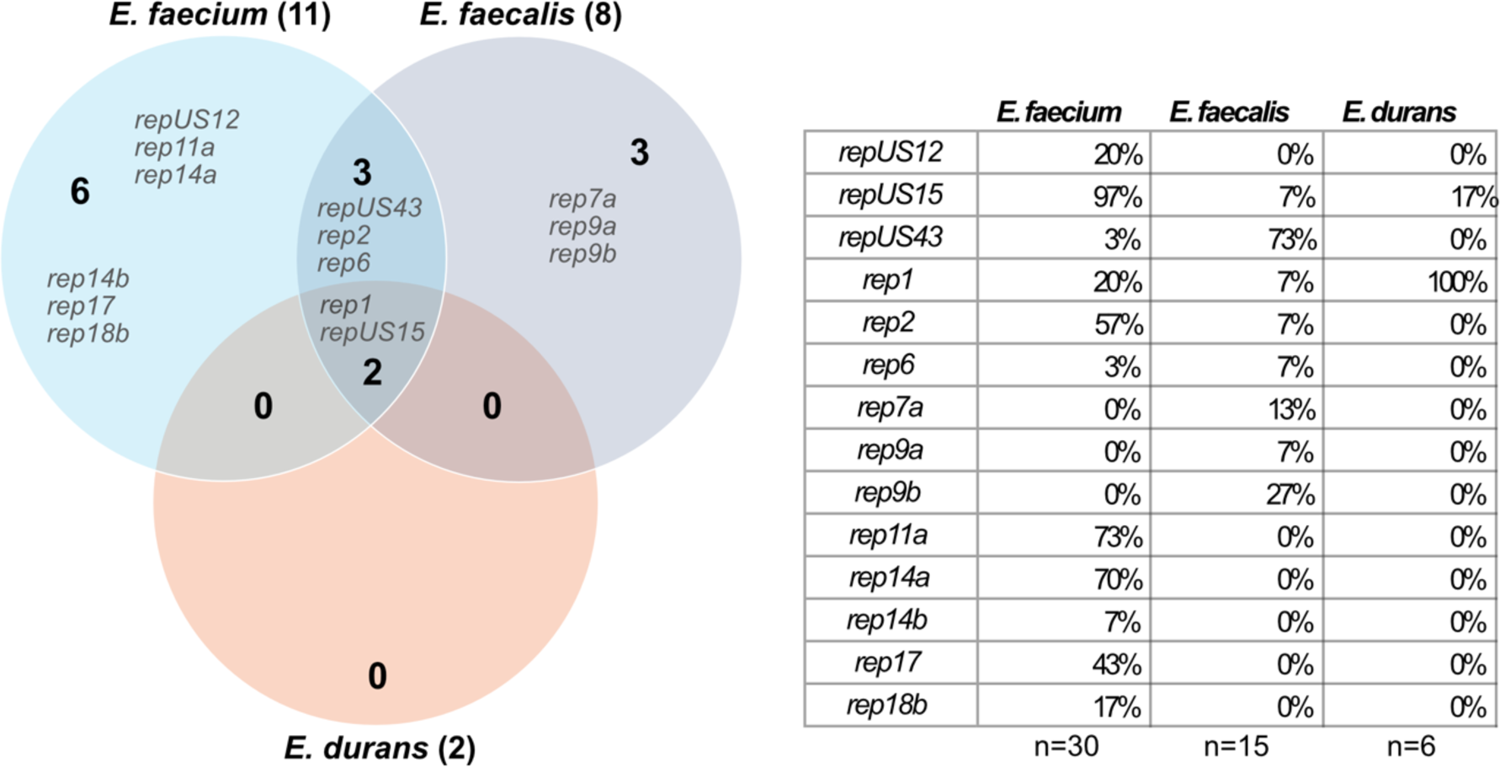
Enterococcal plasmids. Venn diagram illustrating species and indicated plasmids based on PlasmidFinder Rep-screen. The larger bold numbers within the circles represent the total number of unique plasmids carried by the species. The table details the percentages of sequenced isolates found with each given plasmid.

Another interesting observation can be seen in patient HWP030, where a metabolic change occurred between collection days five and six (**Figure 1A**), despite the strains remaining genetically identical (**Figure 1B**). This strain underwent further changes by collection day eight, where 93% of the population (cluster C2-C3, **Figure 1D**) lost its resistance to ciprofloxacin without altering its PhP-type. Interestingly, the replicon identity of rep11a and rep14a fluctuated between 95-100% across the HWP030 collection days (Supplementary Figure 2). Similar plasmid identity variations could be observed in HWP083 and HWP199 (rep14a in both).

Overall, *E. faecium* isolates showed poor metabolic activity for melezitose, inositol and gluconate, but good metabolic activity for L-arabinose, lactose, melibiose, mannitol and sucrose. More varied results were observed for raffinose, sorbitol and amygdalin. For instance, HWP004 and the lone HWP143 isolate did not metabolize sorbitol, the only commonality between them was that these were the only isolates not carrying plasmid 11a. While raffinose metabolism was fairly stable within strains from a single patient, amygdalin metabolism varied greatly even within the same patient. Notably, in HWP004, the smallest subpopulation (cluster 6, 7% subpopulation, **Figure 1C**) during the last collection day had acquired a remarkably strong metabolism of amygdalin. Two additional significant examples are found in patient HWP199, where one isolate uniquely metabolizes mannitol, and in patient HWP143, where one isolate alone metabolizes mannitol, amygdalin and gluconate in the absence of melibiose metabolism. It is noteworthy that all three strains of HWP143 were isolated on the same day, and one of them possessed substantially fewer antimicrobial resistance genes (Supplementary Figure 3).

### Low heterogeneity among *Enterococcus faecalis* isolates

*E. faecalis* formed several minor PhP-type clusters rather than major branches, and with only eight total PhP-types, the isolates were metabolically more similar to each other than what was observed among *E. faecium* isolates. Unlike *E. faecium*, none of the *E. faecalis* isolates were able to metabolize L-arabinose, melibiose, or raffinose similar to e.g. *E. faecium* HWP004 (Figure 3A). Many strains were proficient in metabolizing lactose, melezitose, inositol, sorbitol, mannitol, sucrose, amygdalin and gluconate. Patient HWP003 stood out as the most distinct sample, due to its inability to degrade melezitose. *E. faecalis* from patient HWP167 demonstrated an inability to metabolize inositol (alone in this together with HWP051:L), accompanied with a reduced degradation of gluconate (shared only with HWP051:A). Isolate HWP006:9 exhibited reduced lactose metabolism and was the only isolate for which an ST-type could not be determined. Additionally, HWP006 was the only *E. faecalis* strain carrying plasmids repUS15 and rep1, otherwise only found in our *E. faecium* isolates (Supplementary Figure 2). Notably, patient HWP006 was colonized by *E. faecium* isolates four collection days prior, but they did not carry plasmid rep1; instead, they carried plasmid repUS12 (Figure 1). Due to discrepancies in literature and different databases, it is possible that repUS12 and rep1 are one and the same. Isolate HWP116 was the only isolate showing phenotypic resistance against important antibiotics (Figure 3A), yet it only carried one plasmid shared by most *E. faecalis* (Supplementary Figure 2). Few patients were colonized by *E. faecalis* longitudinally, with only HWP028 serving as an example. The four strains of HWP028 changed their metabolic fingerprint overtime, gradually increasing the metabolic activity of sorbitol, mannitol, sucrose, amygdalin and gluconate. This difference was significant enough to classify HWP028:9A (four collection days to the next collection point) as a separate PhP-type. However, there were no differences in plasmid content, resistance genes (Supplementary Figure 3), or genotype (Figure 3B). The second cluster (C2, Figure 3C) of HWP028 shared the cluster with three other strains, including both isolates of HWP051 (Figure 3C).

**Figure 3.**
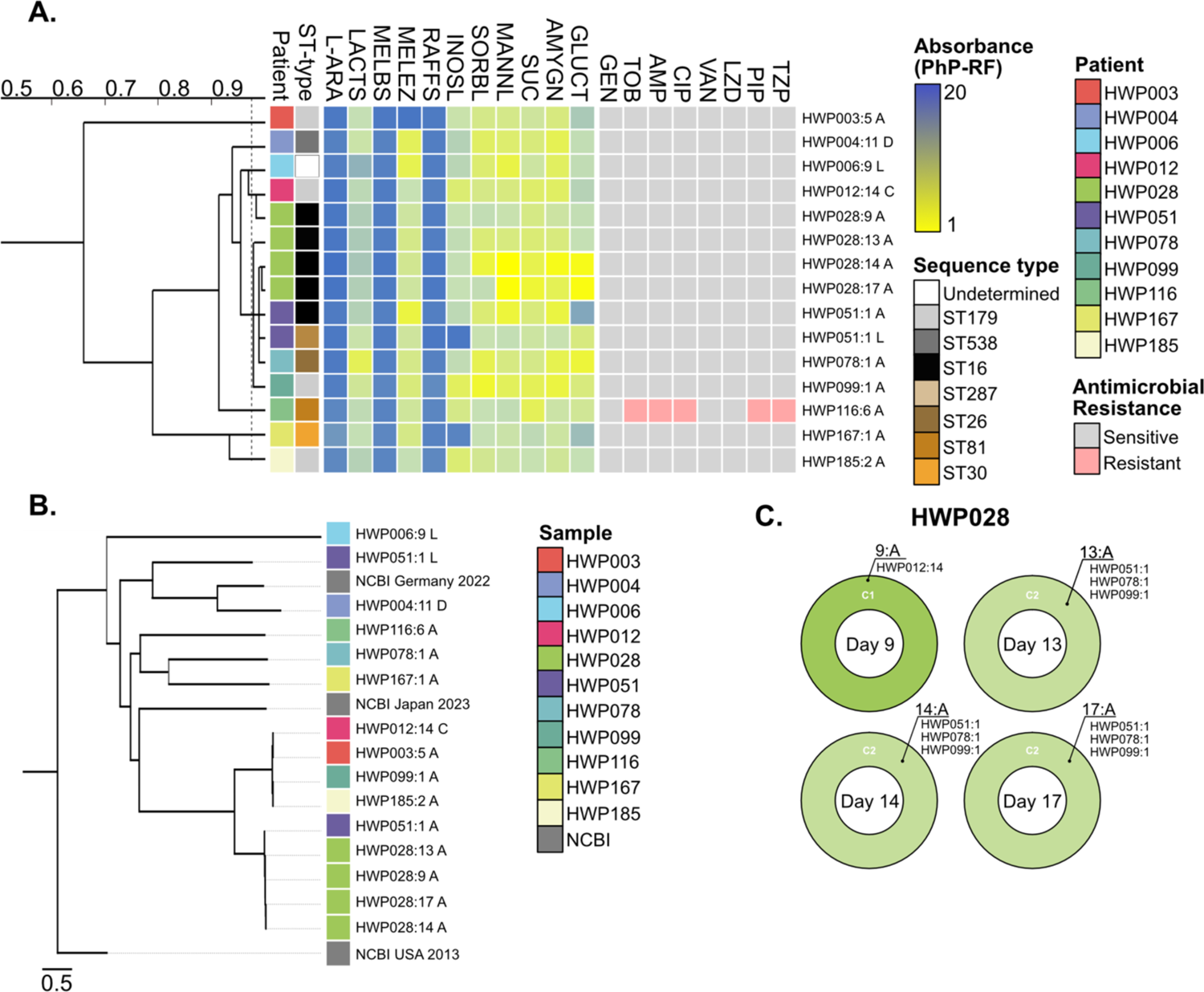
Heterogeneity of *E. faecalis* in the patient cohort. **A.** Dendrogram derived from Unweighted Pair Group Method with Arithmetic Mean (UPGMA) clustering of PhP-RF data (absorbance values at 620nm). The blue-yellow gradient represents a scale from low (blue) to high (yellow) metabolism. Isolates with the most recent branching point above the 0.975 cut-off value (dotted line) are grouped into a PhP-type. Refer to the material and methods or supplementary data for a full list of metabolic reagents. Phenotypic resistance (pink) and susceptibility (gray) was assessed using broth microdilutions according to EUCAST. ST-types were determined based on Illumina WGS. **B.** Phylogenetic tree based on Single Nucleotide Polymorphism (SNP) comparison of Illumina WGS data. Three NCBI strains are included as comparative references based on diversified historical and geographic background. **C.** PhP-type cluster diversity (proportion) based on the number of colonies from the total number screened (n=16) for isolates from patient HWP028.

The two HWP051 strains had different ST-types and grouped differently phylogenetically. Upon closer inspection, HWP051:A (subpopulation 94% of total) carried significantly more AMR genes (Supplementary Figure 3), and a plasmid (repUS43) that HWP051:L did not. In fact, HWP051:L was found with no plasmid at all, a trait shared only with HWP167:1A (Supplementary Figure 2). While it seems likely that this missing plasmid might explain the absence of AMR genes, repUS43 has not been associated with these resistance genes in any other of our strains (Figure 2). This information, coupled with the varied ST-type, indicates a heterogenous population in patient HWP051.

### High heterogeneity was observed among *Enterococcus durans* isolates

For *E. durans*, the sample size was small (two patients), which is also reflected globally with very few strains deposited in NCBI. Overall, *E. durans* shared many metabolic similarities with *E. faecium*, including poor metabolic activity for melezitose and inositol (Figure 4A). None of the strains could metabolize raffinose, similar to *E. faecalis* and some *E. faecium* strains (Supplementary Figure 4). Sorbitol was well metabolized by *E. faecalis* and over half of the *E. faecium* strains, but by none of the *E. durans* isolates. Importantly, all *E. durans* HWP004 strains were isolated on collection day eleven, the same day both *E. faecium* HWP004:E (Figure 1) and *E. faecalis* HWP004:D (Figure 3) were isolated. The *E. durans* isolates almost identically exhibited the phenotype of *E. faecium* HWP004, except for *E. durans* HWP004:11A, which could not metabolize mannitol or sucrose. In terms of AMR, the *E. durans* HWP004 phenotype included resistance against PIP/TZP, similar to *E. faecium*, with one isolate demonstrating resistance towards TOB, just like the as the *E. faecium* strain isolated on the same day. The *E. durans* HWP006 isolates were different from each other, with one isolate capable of metabolizing

**Figure 4.**
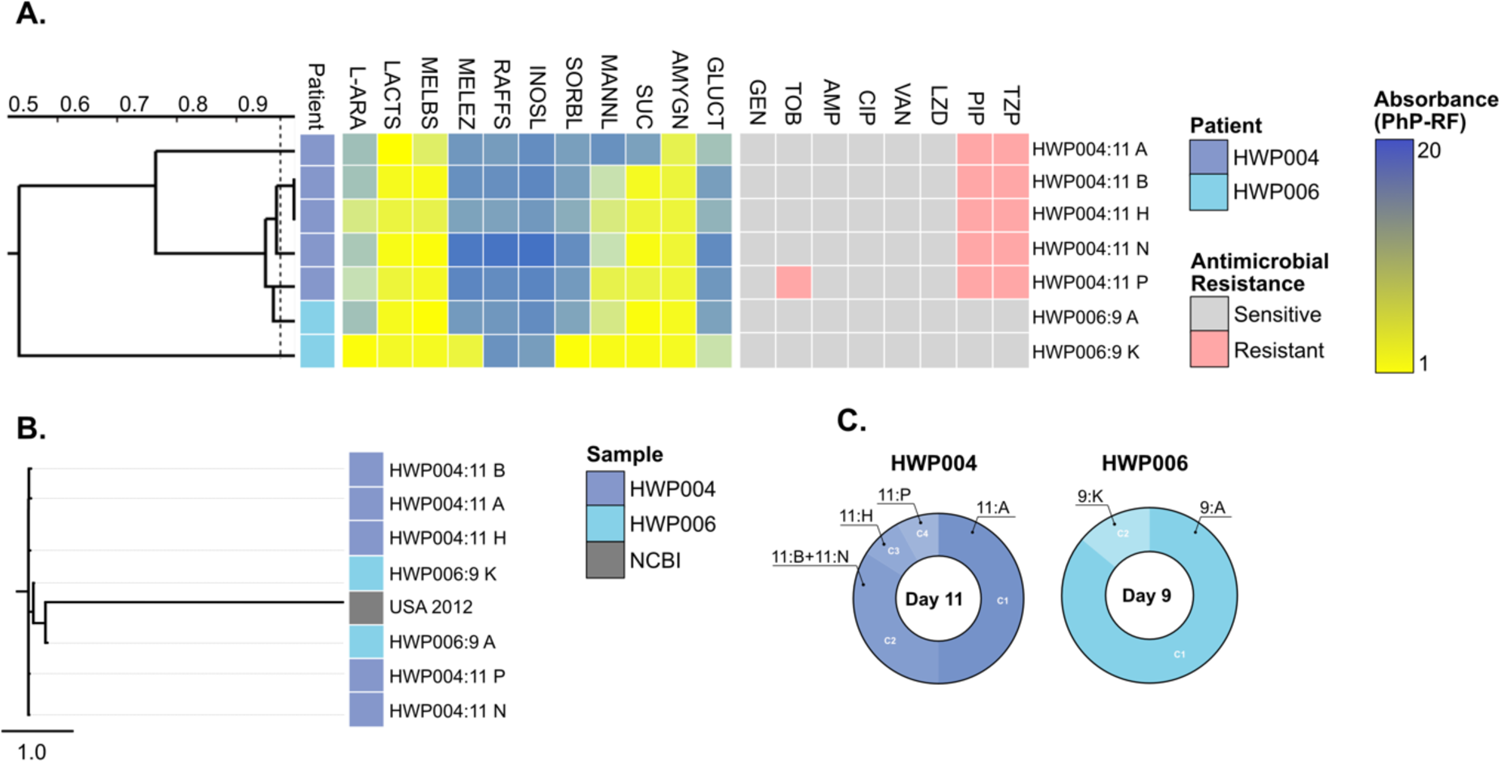
Heterogeneity of *E. durans*. Depiction of *E. durans* diversity in the patient cohort. **A.** Dendrogram derived from Unweighted Pair Group Method with Arithmetic Mean (UPGMA) clustering of PhP-RF data (absorbance values at 620nm). The blue-yellow gradient represents a scale from low (blue) to high (yellow) metabolism. Isolates with the most recent branching point above the 0.975 cut-off value (dotted line) are grouped into a PhP-type. Refer to the material and methods or supplementary data for a full list of metabolic reagents. Phenotypic resistance (pink) and susceptibility (gray) was assessed using broth microdilutions according to EUCAST. **B.** Phylogenetic tree based on Single Nucleotide Polymorphism (SNP) comparison of Illumina WGS data. One NCBI strain is included as comparative reference. **C.** PhP-type cluster diversity (proportion) based on the number of colonies from the total number screened (n=16) for isolates from patient HWP004 and HWP006.

L-arabinose (an *E. faecium* trait) and melezitose (an *E. faecalis* trait). Indeed, this ability might be the reason why HWP006:9K grouped among *E. faecium* isolates instead of *E. durans* isolates (Supplementary Figure 4). None of the HWP006 *E. durans* isolates demonstrated any phenotypic antibiotic resistance, similar to isolate *E. faecalis* HWP006:9 from the same day. Interestingly, *E. durans* HWP006 carried more resistance genes than HWP004, resembling the resistance genotype of *E. faecium* HWP004 (Supplementary Figure 3). Notably, HWP006 had *E. faecium* strains isolated 4-5 collection days prior, and one *E. faecalis* strain isolated on the same day. The *E. faecium*, *E. faecalis* and *E. durans* isolates from HWP006 all carried the repUS15 plasmid, marking the only occurrences of the repUS15 in *E. faecalis* and *E. durans*. Phylogeny indicated that all our strains were similar, but HWP006:9 stood out as the most different isolates (Figure 4B). MLST could not be performed due to insufficient online resources for *E. durans*. Overall, the *E. durans* were diverse and none of the strains survived within the patients for more than one collection day (Figure 4C).

### Hybrid plasmids identified among *Enterococcus* strains

Among the plasmids we identified, two were particularly interesting. The first was from patient HWP004:12, where two plasmid replicons, rep1 and repUS15, were initially identified in *E. faecium* (Figure 2). Following long read assembly, only four contigs were identified: three belonging to the chromosome (total size 2,527,854 bp) and one representing a hybrid plasmid formed from rep1 and repUS15 (112,890 bp) (Figure 5). We subsequently conducted whole genome alignment of this hybrid plasmid contig with two plasmids having the highest similarity according to BLAST (Figure 5). The second notable strain was from patient HWP116:6A. Initially, the repUS43 replicon was identified. After assembly of long read data, only one single contig could be identified (3,001,122 bp). A whole-genome alignment of the plasmid and the bacterial chromosome illustrate that the plasmid appears to have integrated into the chromosome (data not shown).

**Figure 5.**
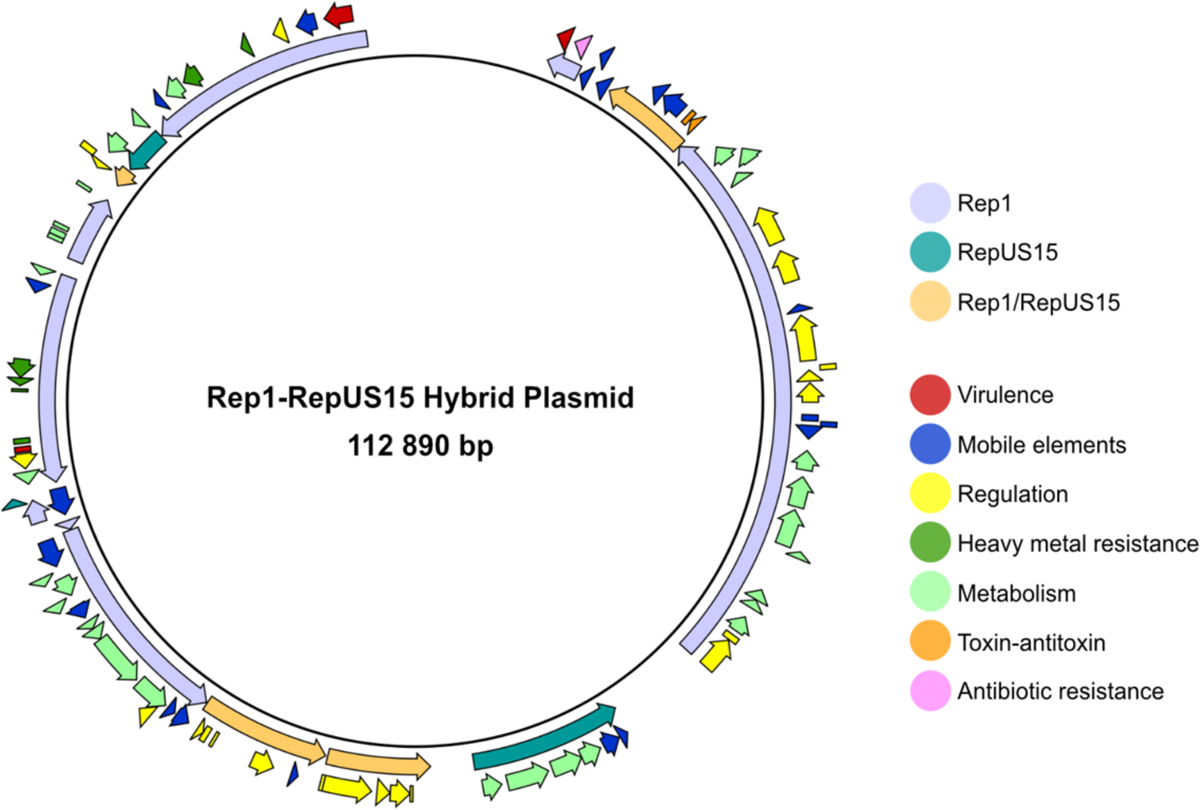
Hybrid plasmid from *E. faecium* HWP004:12B annotated in CLC Genomics Workbench using plasmid sequences from both rep1 and repUS15 plasmids (CP016164 and CP004064). Certain parts of the plasmids are shared and indicated in our plasmid by bright orange (Rep1/RepUS15). Colors indicate suggested functional gene properties based on information received from annotated plasmids.

## Discussion

Differentiating species and clinically relevant strains of *Enterococcus*, often unobserved by conventional screening methods, is challenging and requires a fast and accurate screening scheme. In this study, we used the PhP-RF system, which provides a biochemical fingerprint for various clinical *Enterococcus* species, to investigate the heterogeneity of clinical *Enterococcus* during bladder colonization.

Our study revealed considerable heterogeneity among *E. faecium* isolates. Although underexplored, particularly in the context of bladder infections, it has previously been demonstrated that *E. faecium* infections may involve multiple clones simultaneously (23, 24). Most isolates in our study belonged to the well-characterized clonal complex 17 (CC17, formerly known as clade A1), encompassing strains from ST127 (HWP004) and ST80 (HWP143, HWP199, HWP030, HWP072, HWP083, and HWP095). CC17 is widely recognized as the most significant cluster of *E. faecium*, often linked to enhanced biofilm formation, increased virulence, and AMR in healthcare settings (25, 26). Interestingly, one isolate from HWP143 was identified as ST22 (HWP143:4C), which is not typically associated with pathogenicity (27), but has been proposed as an ancestral lineage of the CC17 cluster (28). Additionally, HWP006 was classified as ST192, a sequence type recently identified as an emerging clone primarily in hospital settings in Germany (29).

Most isolates were classified into distinct PhP-types. HWP143 exhibited clear subpopulations due to the presence of multiple ST-types. In other strains, particularly those from longitudinal samples, observed metabolic variations may suggest metabolic adaptations to the bladder environment. Enterococci are generally thought to lack the necessary enzymes to metabolize urine-based carbon and nitrogen sources, such as creatine and urea (30, 31). However, patients in ICUs with conditions such as diabetes or acute kidney injury (like those in our cohort) may excrete unaltered carbohydrates, including glucose, lactose, and L-arabinose (32). Mannitol is sometimes administered as a diuretic in clinical settings (33), however, this was not the case in our cohort. Isolates from both HWP004 and HWP030 exhibited increased metabolic activity of L-arabinose over time. For HWP143, the majority of the population (81%) belonged to the non-pathogenic ST22, suggesting that the presence of CC17 might have been overlooked by conventional screening methods. In our previous study, we demonstrated that HWP004 was co-colonized by *Candida albicans* on collection day eleven, and by *E*. *coli* on days 12-14 (patient A in cited paper) (5). While most studies have focused on the interaction of *E. faecalis* and other species, the polymicrobial interactions involving *E. faecium* remains less understood.

In patient HWP030, a metabolic shift was observed between collection days five and six, despite genetic similarity, with further alterations leading to a loss of ciprofloxacin resistance in 93% of the population by day eight. Across *E. faecium*, there were variations in AMR within patient isolates, specifically against tobramycin (HWP004, HWP006), piperacillin/tazobactam (HWP004), and ciprofloxacin (HWP030), as well as differences in AMR genes (HWP004, HWP143). Although none of these antibiotics are first-line treatments for *E. faecium*, this finding adds to the growing body of evidence of AMR variability among subpopulations within an infection (34). Indeed, even when the dominant populations were initially resistant, the resistant phenotype rapidly diminished in subsequent collection days, likely due to the absence of selective pressure, at least to a prevalence below our detection threshold.

Plasmid carriage varied among isolates from the same patient, independent of changes in AMR. On collection day 11, the dominant population of HWP004 carried rep1 but lacked repUS15, while a minor subpopulation carried both plasmids. By day 12, Rep1 was absent in two of our strains but reappeared in all strains by day 13, potentially maintained by isolate HWP004:12B. Previous studies have identified rep1 plasmids as conjugative plasmids specific to *E. faecalis*, associated with AMR and virulence, and typically absent in *E. faecium* (**Supplementary Table 2**) (35, 36). Interestingly, all five HWP004:11 *E. durans* isolates carried this plasmid, suggesting horizontal gene transfer within the population. This was not the only instance of rep1 plasmid transfer, as it was also detected in HWP143:4C. Another notable finding was the presence of rep6 in HWP072. Although this plasmid is small and cryptic, it has been proposed as an *E. faecalis*-specific plasmid (35). For isolates HWP030, HWP083, and HWP199, significant nucleotide variations were observed in rep11a (suggested to be a toxin-related plasmid, **Supplementary Table 2**) and rep14a (small and cryptic), as these plasmid replicons showed deviations in identity level between collection days, indicating potential environmental adaptation through plasmid rearrangement.

*E. faecalis* isolates exhibited limited heterogeneity when assessing ST-types within a patient, but demonstrated notable diversity in both ST-type and metabolic profile across different patients. *E. faecalis* ST-types are generally less homogeneous, likely due to their widespread ecological distribution which contributes to the limited understanding of the significance of specific ST-types (37). The predominant ST-type identified was ST179 (HWP003, HWP012, HWP099 and HWP085), which, along with ST16 (HWP028 and one isolate from HWP051) belongs to clonal complex 58 (38). CC58 has been frequently associated with clinical infections, particularly in intensive care units, and with AMR (38, 39). Other identified ST-types included ST538 (HWP004, CC241), ST287 (one isolate from HWP051), ST26 (HWP078), ST81 (HWP116), and ST30 (HWP167), all ST-types of which have previously been linked to clinical infections (40–44). The stand-out example of heterogeneity among *E. faecalis* was observed in patient HWP051, who was colonized by both ST16 (94% subpopulation) and ST287 (6% subpopulation) simultaneously. While prior research on *E. faecalis* heterogeneity has often focused on strain properties, different patient samples, or outbreak scenarios (45–47), the occurrence of mixed enterococcal site infections has not been extensively documented. The rarity of reporting mixed infection might be due to the challenges in distinguishing species, as well as the fact that screenings rarely encompass as many colonies as our current study for the investigation. Indeed, in our previous screen where ten colonies were selected, and the batch later assessed by MALDI-TOF, we did not detect *E. faecalis* HWP004, even though *E. faecalis* had been isolated from blood of the same patient during our urine collection period (5) (patient A in cited paper). This underscores that more pathogenic strains might be missed due the lack of thorough investigations.

In contrast to *E. faecium*, *E. faecalis* isolates did not metabolize L-arabinose, melibiose, or raffinose, but efficiently utilized other carbon sources such as lactose, melezitose, inositol, sorbitol, mannitol, sucrose, amygdalin, and gluconate. These findings align with established knowledge of *E. faecalis* metabolism (48), reinforcing the species-specific metabolic pathways that have been previously documented. Notably, the isolate from patient HWP003 lacked the ability to degrade melezitose, while HWP167 showed deficiencies in metabolizing inositol and gluconate. The isolates from patient HWP051 exhibited distinct metabolic profiles, with key differences in their ability to degrade melezitose, inositol, mannitol, amygdalin, and gluconate. Strains from patient HWP028 demonstrated increased metabolic activity for mannitol, sucrose, amygdalin, and gluconate over time, suggesting the potential importance of these carbon sources during *E. faecalis* bladder colonization. According to our previous investigation, HWP028 had been pre-colonized by *C. albicans* a few days prior to bacterial appearance and was co-colonized by *E. coli* on collection days nine and 13 (patient B in cited paper) (5). Between collection days 9-12, there was no colonization, during which the patient was treated with piperacillin-tazobactam. These species are known to interact and influence gene expression with regards to virulence, but little is yet known about their metabolic dependencies in urine (49, 50).

Most *E. faecalis* isolates harbored the repUS43 plasmid, a conjugative plasmid previously identified in only one of our *E. faecium* isolate. The isolate *E. faecalis* HWP006:9 uniquely carried the repUS15 plasmid, suggesting possible horizontal gene transfer from *E. faecium*. Interestingly, this plasmid was also found in one *E. durans* isolate from the same isolation day, indicating conjugative exchange among all three species present. The repUS15 plasmid has recently been described as a carrier for high-level aminoglycoside resistance and as a potential carrier for *vanA*-mediated vancomycin resistance (51, 52). Tobramycin resistance was only observed in the first *E. faecium* strain carrying the plasmid, and the resistance was lost by the following collection day. The strain HWP078 possessed the highest number of plasmids (six), including one (rep7a) not typically associated with enterococcal plasmids (**Supplementary Table 2**) (35). Additionally, *E. faecalis* HWP116 was the only strain exhibiting phenotypic resistance to tobramycin, ampicillin, ciprofloxacin, piperacillin, and piperacillin-tazobactam, despite carrying few AMR genes and only the marker for the repUS43 plasmid, which turned out to be chromosomally integrated. This case of multidrug-resistant *E. faecalis* is particularly concerning given the general characterization of *E. faecalis* as more virulent but less resistant than *E. faecium* (53). Unlike *E. faecium*, *E. faecalis* plasmids did not exhibit nucleotide changes over time, nor were there longitudinal changes in AMR or AMR genes. Notably, two of the *E. faecalis* strains were not found to carry any plasmid at all, indicating a generally lower frequency of plasmid exchange.

*E. durans* isolates, though less common, shared several metabolic traits with *E. faecium*, such as poor activity for melezitose and inositol and an inability to metabolize raffinose. Unlike *E. faecalis* and many *E. faecium* strains, none of the *E. durans* isolates could metabolize sorbitol. Two *E. durans* strains were initially misidentified as *E. faecium* by MALDI-TOF but were confirmed as *E. durans* through sequencing. This misidentification could only partly be resolved by PhP-RF and is consistent with previous reports highlighting the similarity between *E. durans* and *E. faecium* (54). Despite being prevalent for only one day, these findings are significant as *E. durans* has been associated with higher rates of progression from urinary tract infections to bacteremia (55, 56). Interestingly, likely events of horizontal gene transfer occurred in patients co-colonized with *E. durans* (HWP004 and HWP006). However, these patients also harbored both *E. faecium* and *E. faecalis*, making it impossible to distinguish a single contributing species. As far as we are aware, no studies have been published on horizontal gene transfer or conjugative properties of *E. durans*, making our *E. durans* HWP006:9A one of the first cases of probable interspecies *E. durans* plasmid conjugation. Further studies are needed to prove this transfer.

The *E. durans* strains from patient HWP004, isolated on the same day as *E. faecium* HWP004 and *E. faecalis* HWP004, resembled the *E. faecium* HWP004 phenotype, except for *E. durans* HWP004:11A, which could not metabolize mannitol or sucrose. *E. durans* HWP004 also exhibited a similar antibiotic resistance phenotype to *E. faecium*. In contrast, *E. durans* HWP006 isolates displayed unique metabolic capabilities and carried more resistance genes than HWP004. However, no phenotypic AMR was observed, similar to co-isolated *E. faecalis* HWP006:9. Phylogenetically, strains from HWP004 were similar, while the HWP006 strains stood out both from each other, and from HWP004, underscoring significant heterogeneity among *E. durans* isolates. Notably, all strains carried the rep1 plasmid, which has previously been suggested associated with *E. faecalis* (35), indicating a potential similarity between these two species.

## CONCLUSION

This research provides significant insights into the heterogeneity and metabolic diversity of *Enterococcus* species, particularly *E. faecium* and *E. durans*, across various dimensions. Our findings demonstrate considerable genetic and metabolic diversity among *E. faecium* isolates, with most belonging to clonal complex 17 (CC17), a group known for its enhanced virulence and antibiotic resistance. Unlike earlier studies that focused primarily on single lineages, we identified the presence of multiple lineages and ST-types within individual patients, including ST22, an ancestral lineage of CC17, and ST192, an emerging clone in hospital settings. Notably, metabolic adaptations were observed, such as increased metabolism of L-arabinose, and significant shifts in antibiotic resistance patterns over time. Our work reveals a previously underappreciated metabolic diversity.

*E. faecium* isolates exhibited variability in plasmid content and types. For instance, the rep1 plasmid, typically associated with *E. faecalis*, was found in both *E. faecium* and *E. durans*, suggesting horizontal gene transfer. Variations in plasmid nucleotide sequences over time were also observed, which could reflect environmental adaptation. *E. faecalis* isolates showed less genetic heterogeneity but displayed notable diversity in metabolic profiles across different patients. Evidence of mixed colonization with different ST-types and variations in antibiotic resistance was found. Plasmids such as repUS43 and repUS15, some plasmids not typically associated with *E. faecalis*, were identified, further highlighting the complexity of these infections. *E. durans* isolates, though less common, exhibited metabolic similarities to *E. faecium*. The misidentification of *E. durans* as *E. faecium* by MALDI-TOF, later confirmed through sequencing, underscores the need for accurate identification methods. For the first time, we could demonstrate likely horizontal gene transfer between *E. faecium*, *E. faecalis* and *E. durans*, related to different heterogenous subpopulations within a single patient.

This research reveals the dynamic nature of *Enterococcus* infections, emphasizing significant variability in genetic and metabolic profiles both within individual patients and over time. The study offers novel insights into plasmid diversity and horizontal gene transfer among these species, which may have important implications for our understanding of antibiotic resistance and virulence. These findings underscore the critical need to consider metabolic adaptations and genetic heterogeneity, particularly in clinical settings where conventional screening methods may overlook the presence of multiple clones and lineages, when studying *Enterococcus* infections, as they will come to impact disease severity and treatment outcome.

## MATERIAL AND METHODS

### Sample collection

Urine sample collection was done as previously described (5). Briefly, urine was collected from the catheter into sterile vacutainer tubes and transported cold. All urine samples were stored in cryovials with 10% dimethyl sulfoxide (DMSO) at −80°C. The same study performed a full screen of all urine samples and collection days from 210 patients. This current study focused on *Enterococcus*-positive urine samples (n=29) from the initial screen, from which 16 colonies per urine sample (n=464) were further investigated.

### *Enterococcus* identification and isolation

For *Enterococcus* identification, 100µl of thawed urine was streaked on Brilliance™ UTI Clarity™ agar (UTI agar) plates, incubated for 24 hours at 37°C. Urine was diluted in phosphate-buffered saline (PBS) to meet *Enterococcus* colony-forming units (CFU) around 50-100 CFU per plate. To investigate heterogeneity, 16 *Enterococcus* colonies were picked from each plate and re-streaked on Brain Heart Infusion (BHI) agar, incubated at 37°C for 24 hours. If the number of single *Enterococcus* colonies was less than 16, or the urine contained polymicrobial colonization complicating collection, additional streaking on UTI agar was performed. This step was to ensure the isolates were pure *Enterococcus* colonies.

### PhP-RF system screening

The PhP-RF system is a rapid phenotypic screening method from the PhenePlate™ typing system assessing the kinetics of biochemical reactions involved in prokaryotic metabolism. It is composed of a 96-well microtiter plate with eleven different dehydrated substrates coated at the bottom (**Supplementary table 1**). In our study, all 16 isolates per urine sample were tested using the PhP-RF system. After three readings of absorbance, the mean value was calculated to compare similarities. Pairwise similarities between isolates showed their biochemical fingerprints and were calculated as correlation coefficients (21). These correlation coefficients were used by the PhPWIN software to create a similarity matrix, later clustered into a dendrogram using Unweighted Pair Group Method with Arithmetic Mean (UPGMA). Isolates sharing biochemical fingerprint similarity below 0.975 were recognized as different PhP-types indicating metabolic heterogeneity.

A suspending medium containing 0.011% bromothymol blue, 0.2% proteose peptone, 0.05% yeast extract, 0.5% sodium chloride, and a 0.2M solution of phosphate buffer (Na_2_HPO_4_ + NaH_2_PO_4_) at pH 7.5 was prepared according to the instruction manual of the PhP-RF system (18). *Enterococcus* colonies from BHI agar were picked with sterile toothpicks and added to the first column of the PhP-RF plate, and then aliquoted from column one to each of the reagent wells in the consecutive columns. The inoculated PhP-RF plates were incubated inside a moist microaerophilic chamber for 64 hours at 37°C. Absorbance at 620 nm was measured after 16, 40, and 64 hours with a Spark^®^ 10M Multimode Microplate Reader (Tecan Nordic AB). The tested compounds included L-arabinose (L-ARA), lactose (LACTS), melibiose (MELBS), melezitose (MELEZ), raffinose (RAFFS), inositol (INOSL), sorbitol (SORBL), mannitol (MANNL), sucrose (SUC), amygdalin (AMYGN) and gluconate (GLUCT).

### Antimicrobial resistance testing

The antimicrobial resistance (AMR) tests were carried out in round-bottom 96-well plates. Concentrations were only tested at clinical breakpoint provided by EUCAST (v 14.0)(57). Bacterial suspensions were calibrated in 0.9% NaCl to 0.5 McFarland standard units, diluted in Mueller Hinton (MH) broth, and added to each well at a final concentration of 5×10^5^ cells/ml. Positive control wells had no antibiotics (growth control) and negative controls wells had no bacteria (media control). Both positive and negative control wells were included in every plate.

Antibiotics used included gentamicin (GEN), tobramycin (TOB), ampicillin (AMP), ciprofloxacin (CIP), vancomycin (VAN), linezolid (LZD), piperacillin (PIP) and piperacillin-tazobactam (TZP). Tazobactam concentration was added as 4mg/L for all TZP wells. The plates were incubated for 18-20 hours at 37°C.

### Strain identification and WGS

Minimum one representative isolate per PhP-type was used for strain identification. Fresh colonies on BHI plates were identified with matrix-assisted laser desorption/ionization time-of-flight mass spectrometry (MALDI-TOF, Bruker Biotyper^®^), additionally confirmed by whole-genome sequencing (WGS). The same isolates used for MALDI-TOF were suspended in BHI broth and prepared as overnight cultures in arrangement for DNA extraction (*E. faecium*: n=31, *E. faecalis*: n=18, *E. durans*: n=5). DNA extraction was performed on overnight cultures using Lucigen MasterPure™ Complete DNA and RNA Purification Kit (Cat. No. MC85200). DNA quantity was checked with a Qubit 2.0 Fluorometer for broad rage (BR) double stranded DNA. Extracted DNA was sent to BMKGENE for Illumina-based short read whole genome sequencing. For a subset of strains, we carried out complementary Oxford Nanopore long-read sequencing, later corrected using the Illumina reads.

### Sequence analysis

Sequence analysis was performed in CLC Genomics Workbench v24.0.1, as described from our lab (58). Selected isolates belonging to different PhP-types were sequenced by Illumina, and reads were paired, trimmed, and assembled into contigs. Species were confirmed using NCBI nucleotide Basic Local Alignment Search Tool (BLAST) on the largest contigs assembled. To compare isolates genetically, we crafted a phylogenetic tree using single nucleotide polymorphism (SNP) inferred phylogeny from CSI Phylogeny 1.4, standard settings (59). Phylogenetic trees were generated per species. Sequence types (ST) of our strains were determined using Multi-Locus Sequence Typing on *de novo* assembled genomes (MLST 2.0) (19, 60–64). ResFinder v4.1 was used to identify AMR genes based on unassembled reads (65), and plasmid finder (v2.1) was used as an indication for the presence of plasmids (66). A subset of strains with plasmids were Nanopore sequenced and assembled (Long Read Support v24.0), and confirmed plasmids were aligned (Whole Genome Alignment V24.0) to reference plasmids (NCBI, BLAST) in CLC Genomics Workbench.

### Visualization

PhP-type trees were generated with the PhenePlate™ Software while phylogenetic SNP trees were generated with CSI Phylogeny 1.4. AMR profile illustrations were created using Microsoft^®^ Excel (v. 16.86). Illustrations for PhP-types within a patient (circle diagrams) were created using GraphPad Prism (v. 10.2.3). Venn diagram and combination of data sets into single figures were performed using Affinity Designer (Serif/Affinity, v. 1.10.0). Hybrid plasmid was constructed and visualized in CLC Genomics Workbench.

## ETHICS STATEMENT

Data found in this research are part of the PronMed study approved by the National Ethical Review Agency [Dnr 2017/043 (with amendments 2020-01623, 2020-02719, 2020-05730, and 2021-01469) and 2022-00526-01)] and listed at ClinicalTrials.gov (NCT03720860). Informed consent was obtained from the patient or next of kin. The Declaration of Helsinki and its subsequent revisions were followed. Adult ICU patients admitted between February 25^th^ 2021 and September 3^rd^ 2021 with reverse-transcription polymerase chain reaction (RT-PCR) positive nasopharyngeal swabs were prospectively recruited to the study. Exclusion criteria were pregnancy, current breastfeeding, and age under 18. End of follow-up was defined to the end of ICU treatment.

## AUTHOR CONTRIBUTIONS

TZ, PK, EJ and HW: conceptualization. PK, TZ: writing - original draft. TZ, PK: data curation. TZ, PK: formal analysis. HW, JJ, EJ: funding acquisition. PK, TZ, HW: investigation. TZ, PK, HW, EJ: methodology. PK, HW: project administration. PK, HW, EJ: validation. PK, TZ, HW, EJ, JJ: writing - review and editing. All authors contributed to the interpretation of results and critical review of the manuscript.

## ACKNOWLEDGMENTS

We would like to thank Andreas Wallberg, Carl-Johan Rubin and Marie Wrande for their contributions in Nanopore sequencing.

## DATA AVAILABILITY

Not applicable.

## SUPPLEMENTARY

**Supplementary Figure 1.**
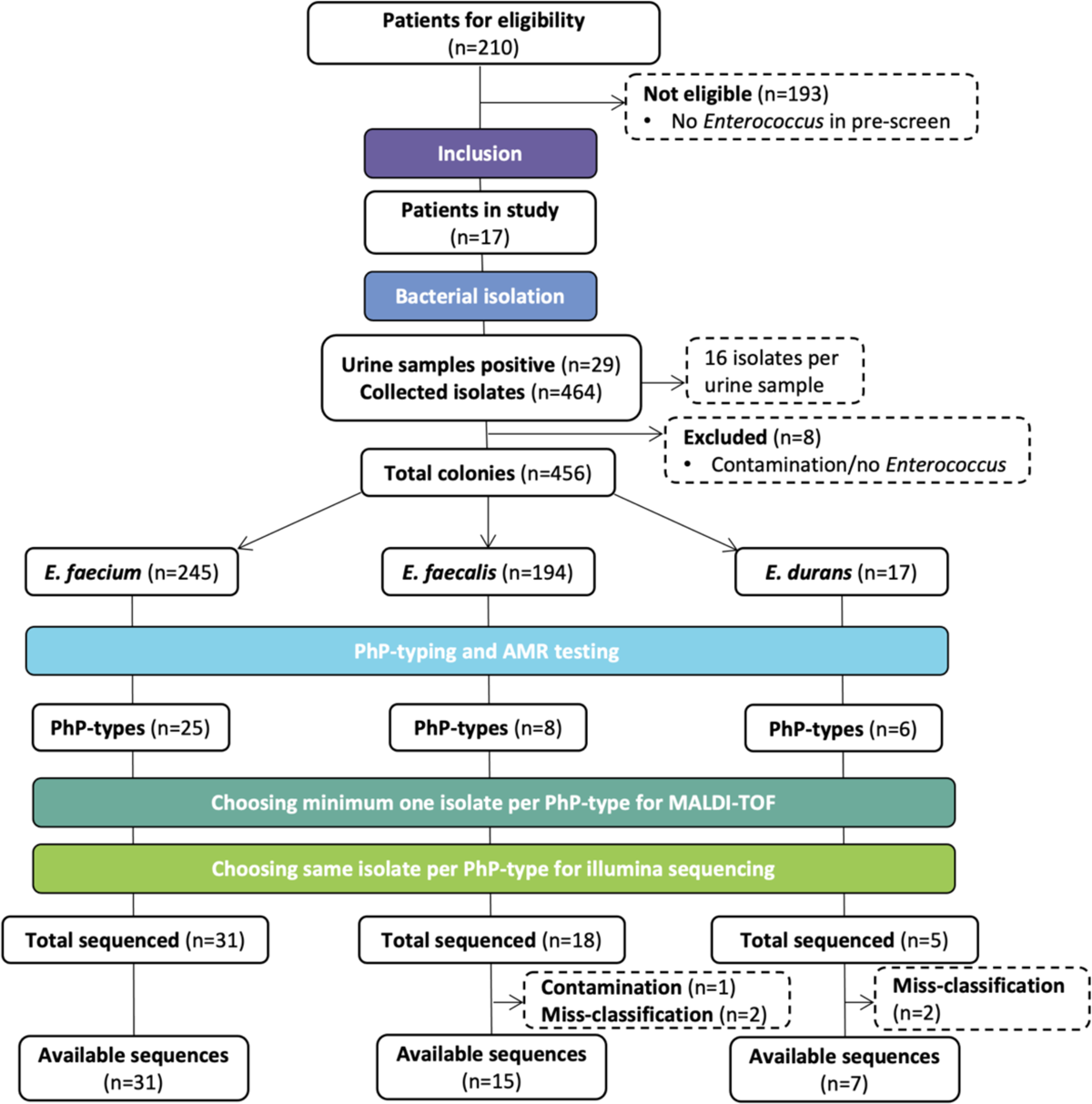
Illustrative pipeline of sample processing from patient eligibility to final count of included sequences.

**Supplementary Figure 2.**
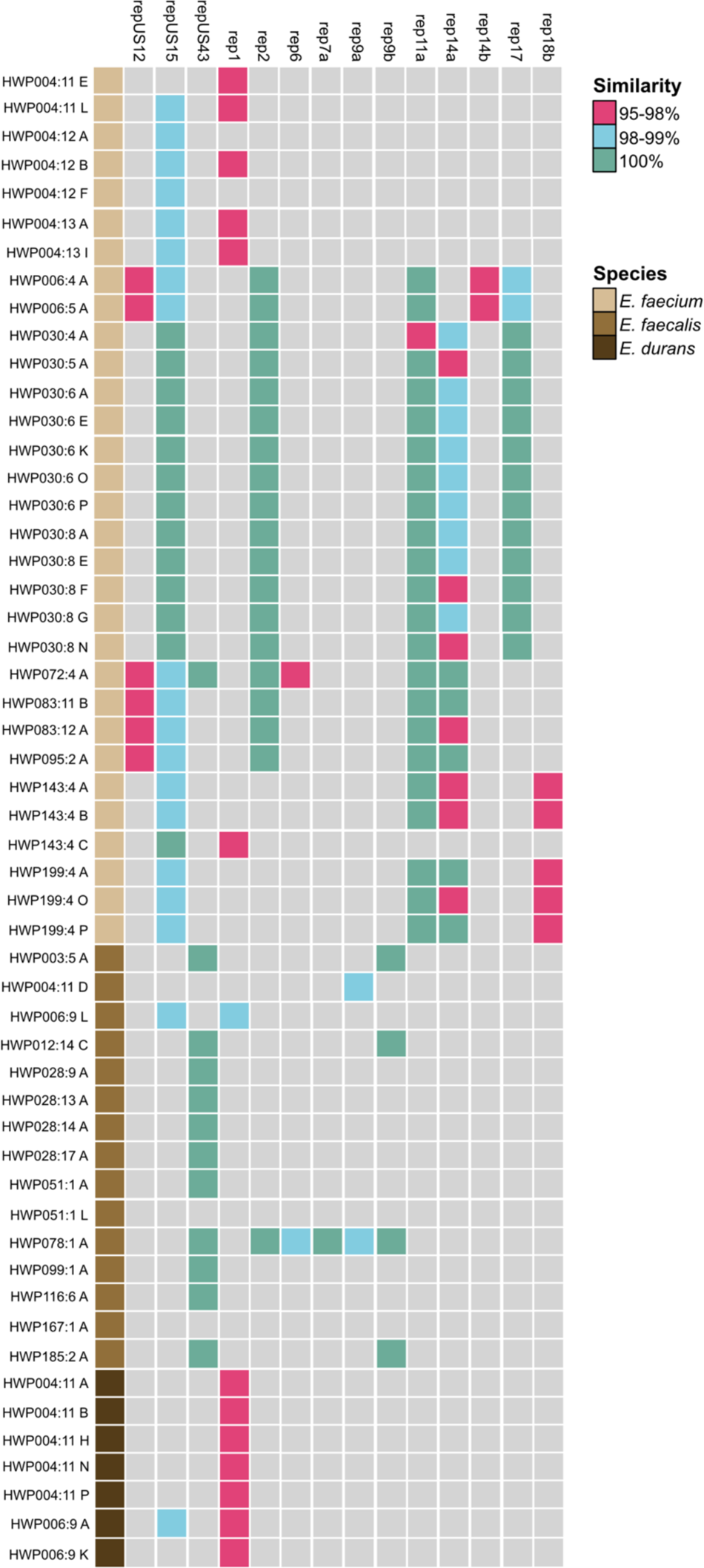
Illustration of plasmid content as identified by PlasmidFinder. Color indicates presence (red, blue, green) and no presence (gray) of plasmid. Species and strains (left) and plasmid replicons (top) are indicated.

**Supplementary Figure 3.**
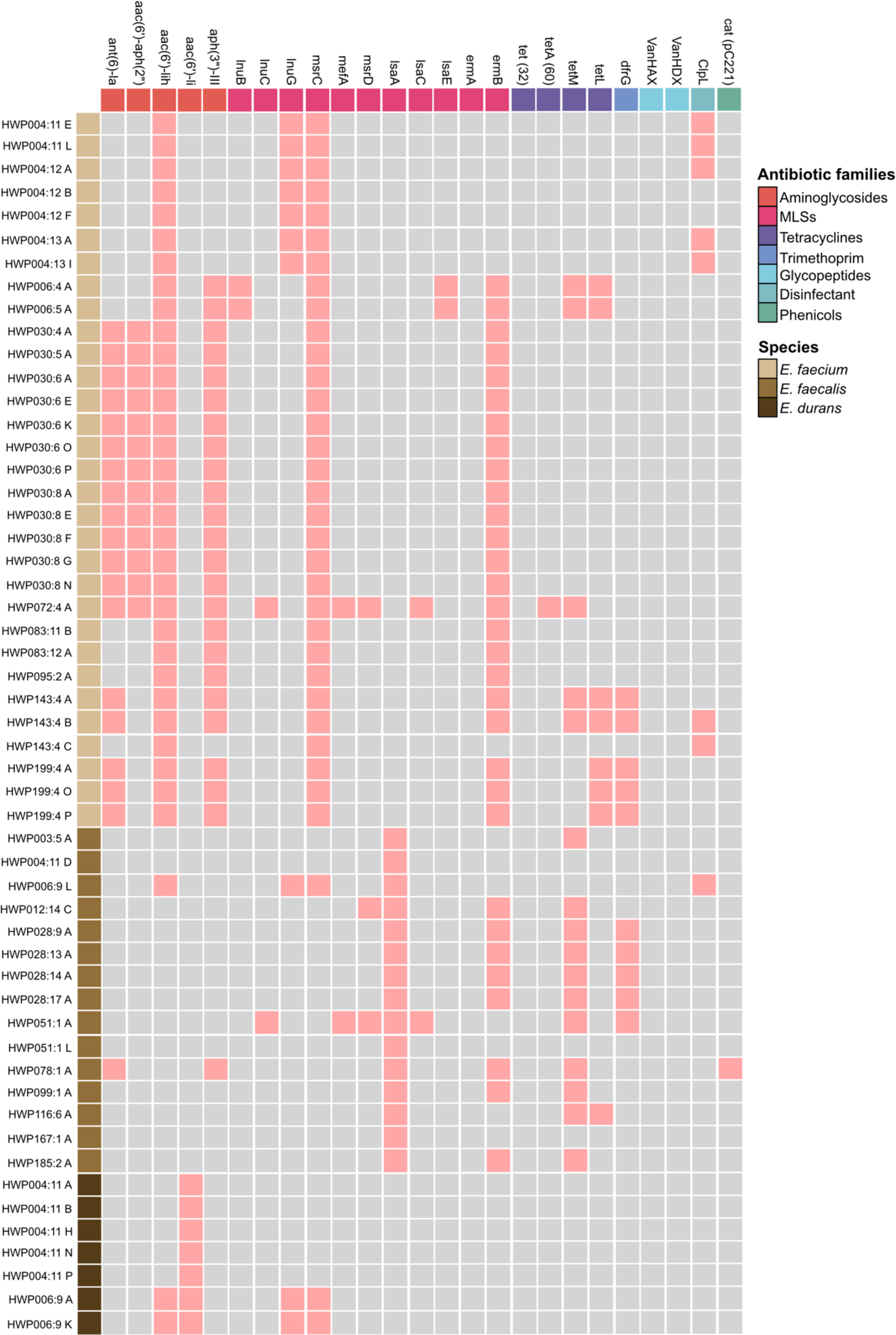
Illustration of antibiotic resistance genes as identified by ResFinder. Color indicates presence (pink) and no presence (gray) of the corresponding gene. Species and strains (left) and resistance gene with the corresponding antibiotic family grouping (top) are indicated.

**Supplementary Figure 4.**
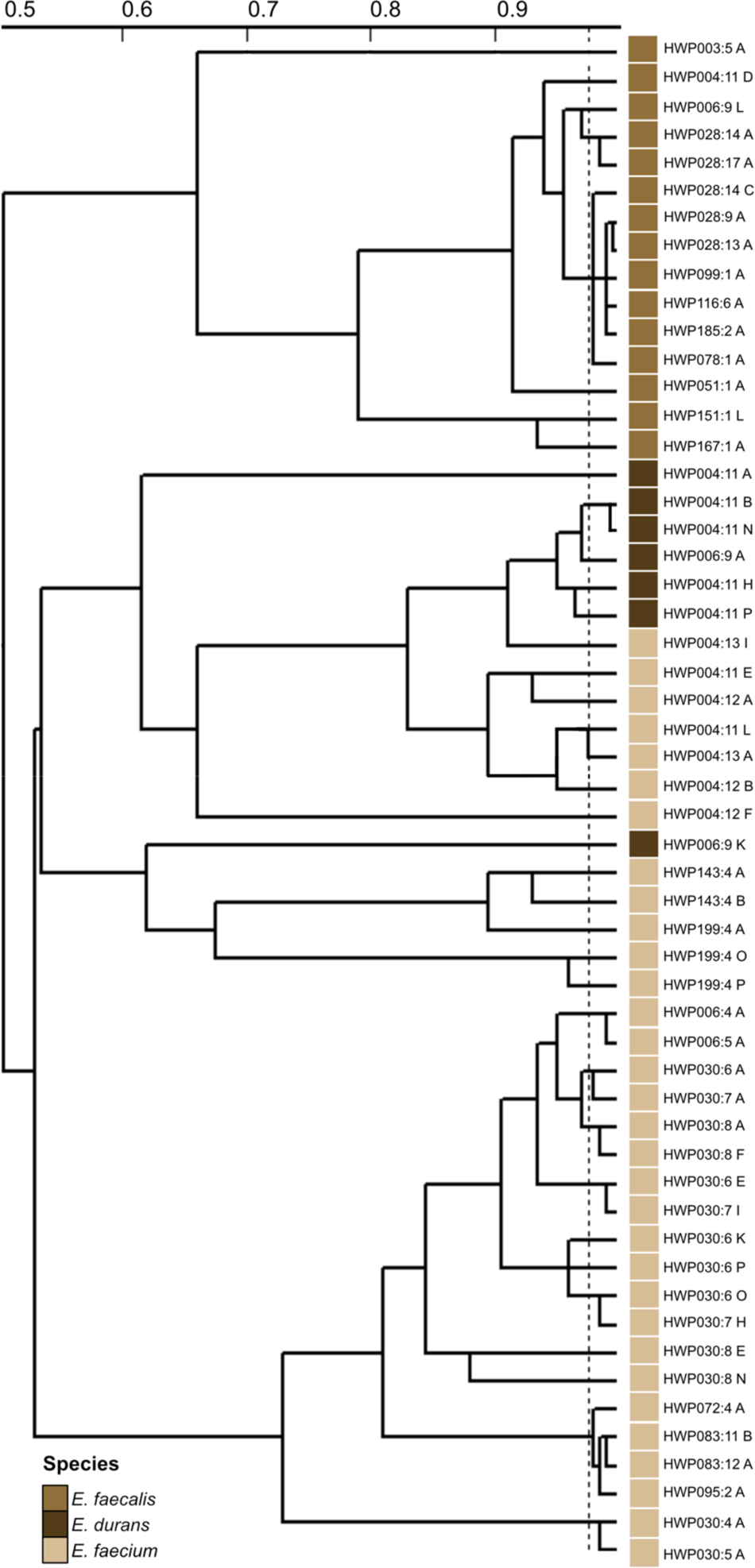
PhP types as previously presented. All species combined into one PhP tree for interspecies comparison.

---

**Supplementary table 1.** Carbon sources.

|  |  |  |
| --- | --- | --- |
| L-arabinose | L-ARA | Monosaccharide found in e.g. dairy |
| Lactose | LACT | Disaccharide found in e.g. dairy |
| Melibiose | MELBS | Disaccharide found in e.g. legumes, byproduct of raffinose |
| Melezitose | MELEZ | Trisaccharide found in e.g. tree sap, tree fungus or honeydew |
| Raffinose | RAFFS | Trisaccharide found in e.g. plant tissue |
| Inositol | INOSL | Sugar alcohol found in e.g. whole grain, fruits and nuts |
| Sorbitol | SORBL | Sugar alcohol found in e.g. fruits, sugar-free products and processed food |
| Mannitol | MANNL | Sugar alcohol found in e.g. mushrooms, vegetables, sugar-free products and processed food |
| Sucrose | SUC | Disaccharide found in e.g. sugarcane and fruits |
| Amygdalin | AMYGN | Cyanogenic glycoside found in the e.g. seeds of certain fruits |
| Gluconate | GLUCT | Conjugate base of organic base found in berries and fermented food |

**Supplementary Table 2.** Enterococcal plasmids Data in this table is based on previous studies (35, 36, 67, 68).

| Replicon | Alternative name | Hosts | Characteristic | Host range |
| --- | --- | --- | --- | --- |
| repUS12 | Rep1 | - | - | - |
| repUS15 | RepA_N | - | Conjugative<br>T4SS | Narrow |
| repUS43 | Rep_trans | - | Conjugative<br>T4SS | - |
| rep1 | Inc18 | *E. faecalis*<br>Other G+ | Toxins<br>AMR<br>Conjugative | Broad |
| rep2 | Inc18 | Enterococci | Toxins<br>AMR<br>Conjugative | Narrow |
| rep6 |  | *E. faecalis*<br>Other G+ | Small, cryptic<br>Rolling circle | Broad |
| rep7a | pT181 Class 1 | Other G+ | AMR<br>Small | Broad |
| rep9a | RepA_N | *E. faecalis* | Sex-pheromone<br>AMR<br>Virulence | Narrow |
| rep9b | RepA_N | *E. faecalis* | Sex-pheromone<br>AMR<br>Virulence | Narrow |
| rep11a | Rep3 | Enterococci | Toxins | Narrow |
| rep14a | Rep_trans | *E. faecium* | Conjugative<br>Small, cryptic | Narrow |
| rep14b | Rep_trans | *E. faecium* | Conjugative<br>Small, cryptic | Narrow |
| rep17 | RepA_N | *E. faecium* | Conjugative<br>AMR | Narrow |
| rep18b | Rep3 | Enterococci | Unknown | Narrow |

